# Caribbean multi-centre study of *Klebsiella pneumoniae*: whole genome sequencing, antimicrobial resistance and virulence factors

**DOI:** 10.1101/541136

**Authors:** Eva Heinz, Richard Brindle, Andrina Morgan-McCalla, Keisha Peters, Nicholas R. Thomson

## Abstract

The surveillance of antimicrobial resistant isolates has proven to be one of the most valuable tools to understand the global rise of multidrug-resistant bacterial pathogens. We report the first insights into the current situation in the Caribbean, where a pilot project to monitor antimicrobial resistance through phenotypic resistance measurements combined with whole-genome sequencing was set up in collaboration with the Caribbean Public Health Agency (CARPHA). Our first study focused on *Klebsiella pneumoniae*, a highly relevant organism amongst the Gram-negative opportunistic pathogens world-wide today causing hospital, as well as community-acquired, infections. Our results show that not only carbapenem resistance, but also hypervirulent strains, are circulating in patients in the Caribbean. Our current data does not allow us to infer their prevalence in the population. We argue for the urgent need to further support antimicrobial resistance surveillance and stewardship in this almost uncharted territory, which can make a significant impact on the reduction of antimicrobial usage.

**DATA SUMMARY:**

-Raw sequencing data is deposited at the sequence read archive (SRA), and assemblies are deposited at GenBank, accession numbers for all are given in Dataset S1.
-The data of measured resistance phenotypes (Vitek) is also provided in Dataset S1.
-The tree file and associated metadata can be investigated and downloaded through the free online platform microreact (https://microreact.org/project/S1-a7KAkV).
✓ **I/We confirm all supporting data, code and protocols have been provided within the article or through supplementary data files. ⊠**

**Significance as a BioResource to the community:** This BioResource contains whole-genome sequence data of 270 *Klebsiella pneumoniae* isolates, information about encoded resistance genes and the phylogeny of the isolates, their distribution in the global *K. pneumoniae* population, and their resistance phenotype data as determined by the VITEK 2 compact system. The isolates are recent (2017 through 2018) and represent clinically relevant patient isolates from 15 different sites in 12 Caribbean states. These data will be of interest for researchers working on *K. pneumoniae* and other opportunistic pathogens as well as those interested in mobile genetic elements carrying antimicrobial resistance cassettes. Our data is the only recent survey of antimicrobial resistant opportunistic pathogens from multiple sites within the Caribbean, and is of high significance for the global surveillance of *K. pneumoniae* and antimicrobial resistance elements. This BioResource is made available through data tables provided with this article, as well deposition of the raw data in the relevant archives, and an interactive platform (microreact) to enquire and download analyses (phylogenetic tree, metadata).

## INTRODUCTION

The increasing level of antimicrobial resistance (AMR) in bacterial pathogens is one of the biggest worldwide threats for public health [1]. The spread is amplified as mobile resistance elements can cross both geographic and species borders, and especially *Enterobacteriaceae* are prone to disseminating plasmids encoding antimicrobial resistance genes [2]. Monitoring the spread of resistant strains and resistance elements is further complicated as most of these bacteria are opportunistic pathogens which can be carried asymptomatically as part of the human microbiota, and the mobility of people today thus greatly contributes to their world-wide spread. The phenomenon has been recognised by the major public health agencies, and several surveillance programs are set in place to assess the prevalence of AMR in bacteria. This facilitates more informed decisions for interventions, guidelines for AMR practice and contributes to our understanding of the mechanisms leading to dissemination of AMR and the emergence of new resistances or high-risk lineages [1].

The Caribbean is a setting with a highly mobile population. The Caribbean Public Health Agency (CARPHA) incorporates 21 island states and 3 located in the Central and South American mainland (http://carpha.org/Who-We-Are/Member-States). This project was launched as part of a longitudinal antimicrobial resistance surveillance strategy in the Caribbean, initiated with funding from the United States Centre for Disease Control and Prevention (CDC) in 2016, to provide insight into the current state of AMR resistance and to develop an antimicrobial stewardship programme. Antimicrobial resistance surveillance is essential to identify potentially problematic clones and resistances, prevent future and recognise on-going epidemics, set measures to prevent further spread of high-risk clones, and better inform antimicrobial usage for health care workers. To be effective, antimicrobial resistance surveillance needs to be established in combination with infection control and antimicrobial stewardship.

Our pilot project targeted *Klebsiella pneumoniae*, a member of the *Enterobacteriaceae* and recognised as one of the greatest threats for public health amongst multi-resistant Gram-negative opportunistic pathogens [3][4]. A Caribbean-wide point prevalence survey showed high usage of beta-lactam antibiotics, especially third-generation cephalosporins, as well as quinolones, macrolides and a considerable degree of carbapenem usage (Figure 1). We report the results of the first two surveys of isolates collected across the CARPHA member states (CMS). The first batch of isolates were collected in early 2017 with a second set of samples submitted to CARPHA during first half of 2018. Unfortunately, funding for this project has ceased along with CARPHA-based AMR surveillance. We provide an important snapshot of AMR across the Caribbean, including phenotypic and genomic data, analysis of the virulence and antimicrobial determinants, and the phylogenetic distribution of the Caribbean isolates in the context of the global *K. pneumoniae* population structure [5].

**Figure 1:**
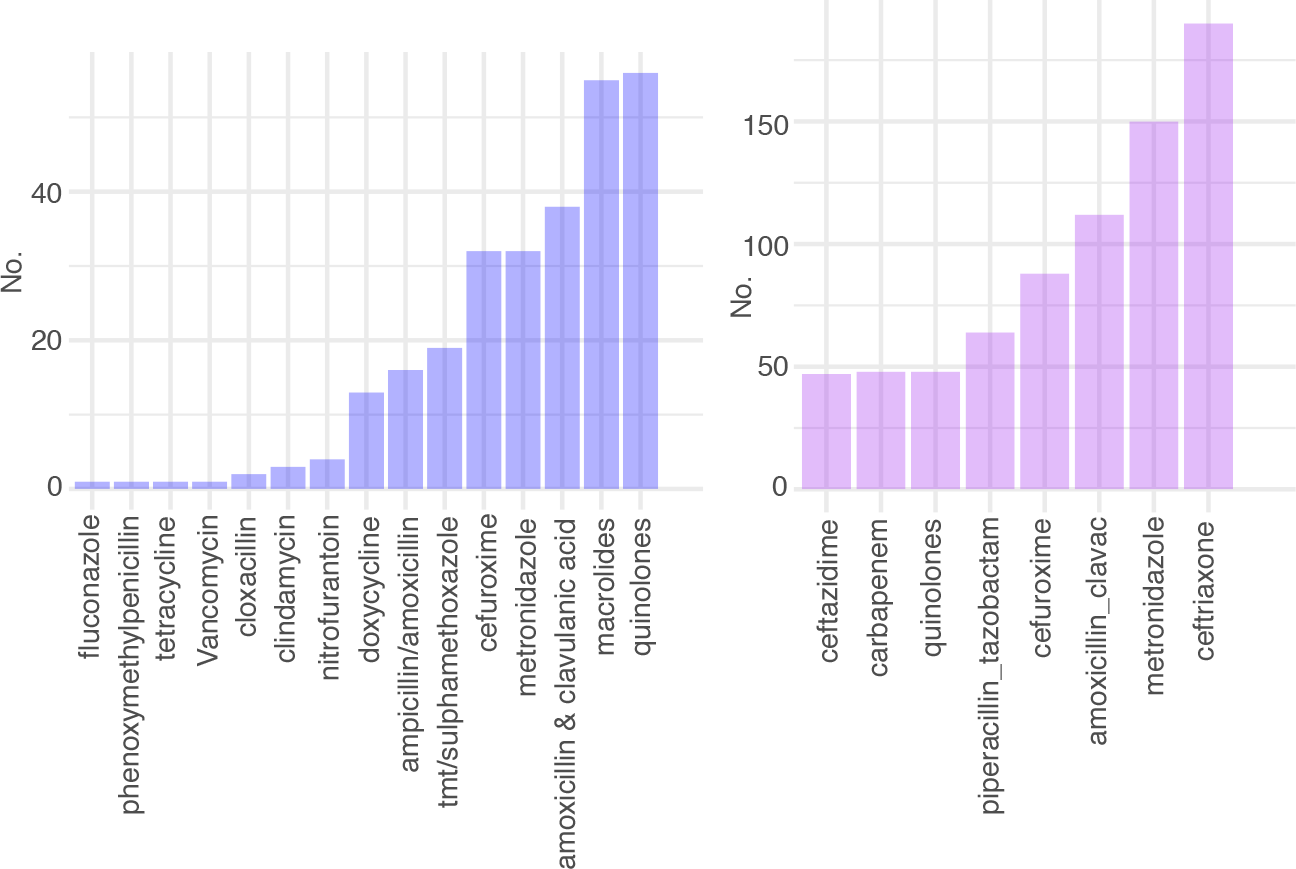
Antimicrobial usage in the Caribbean. From the 2018 point prevalence survey of antimicrobial usage. Shown are the main inpatient prescribed classes for oral (left panel) and intravenous (right panel) antimicrobials in selected hospitals.

## METHODS

### Antimicrobial usage data

The point prevalence (PPS) data was collected, as part of the Caribbean antimicrobial stewardship training programme, between March and May 2018. A streamlined version of the WHO data collection form was used and pharmacist teams collected data from hospital wards and clinics. Thirteen hospitals from 10 states submitted data in time for analysis in June 2018. A total of 1248 patients were reviewed of which 681 patients had been prescribed 1136 antibiotics.

### Sample collection

Isolates were submitted by hospitals from the CARPHA members states; Antigua, Barbados, Belize, Bermuda, Cayman Islands, Dominica, Grenada, Haiti, Saint Kitts, Saint Lucia, Saint Vincent, and Trinidad. Contributing CMS are encoded to anonymise the hospitals and states. The isolates were not selected for submission in a formal or structured fashion and submission was dependent on the availability of transport media and staff availability. The isolates were mainly from bloodstream, wounds and urine samples, but also from a wide range of other sources, including cerebrospinal fluid; further details on the specimens as well as all accession numbers and sequencing details are given in Table S1. Phenotypes and antimicrobial susceptibilities were determined using the VITEK 2 compact system (bioMerieux Inc., 100 Rodolph St., Durham, NC27712, USA) within the microbiology laboratory within CARPHA, Port of Spain, Trinidad.

### Sequencing and analysis

DNA was isolated using the QIAamp^®^ DNA Mini kit following manufacturer’s instructions within the CARPHA laboratory, Illumina sequencing libraries with a 450-bp insert size were prepared according to the manufacturer’s protocols and sequenced on an Illumina HiSeq2000 with paired-end reads with a length of 100bp; accession numbers of all samples are given in table S1. The data was de novo assembled using the pipeline as described in [6], and annotated with prokka [7]. Multiple locus sequence types were predicted as described previously [8]. Presence of antimicrobial resistance and virulence genes were investigated using kleborate (https://github.com/katholt/Kleborate). The core gene alignments were generated using roary v3.7.0 [9] with the default conditions using mafft v7.205 [10]; SNPs were first extracted using snp-sites v2.3.2 [11], and then a maximum likelihood tree was calculated with RAxML v8.2.8 [12]. The data was visualized with the ggtree and ggplot2 packages in R [13][14].

## RESULTS

The isolates included in this study were submitted by a total of 15 different hospitals in 12 CMS (Figure 2A). Although the majority were isolated from primary urine or blood samples, other sample sites included wound infections or invasive isolates causing (liver) abscess, cerebrospinal fluid (CSF) isolates and one case of meningitis (Figure 2B). A significant proportion of the isolates included in this study were resistant to all commonly used classes of antimicrobials, with the exception of the carbapenems and tigecycline. Phenotypic screening confirmed a high level of extended-spectrum beta-lactamase resistance and, to a similar extent, reduced susceptibility to other antimicrobial classes such as aminoglycosides and fluoroquinolones (Figure 2C); whereas only a small proportion of the isolates were phenotypically carbapenem resistant. We also note a relatively low proportion of amikacin and piperacillin-tazobactam resistance (Figure 2C).

**Figure 2:**
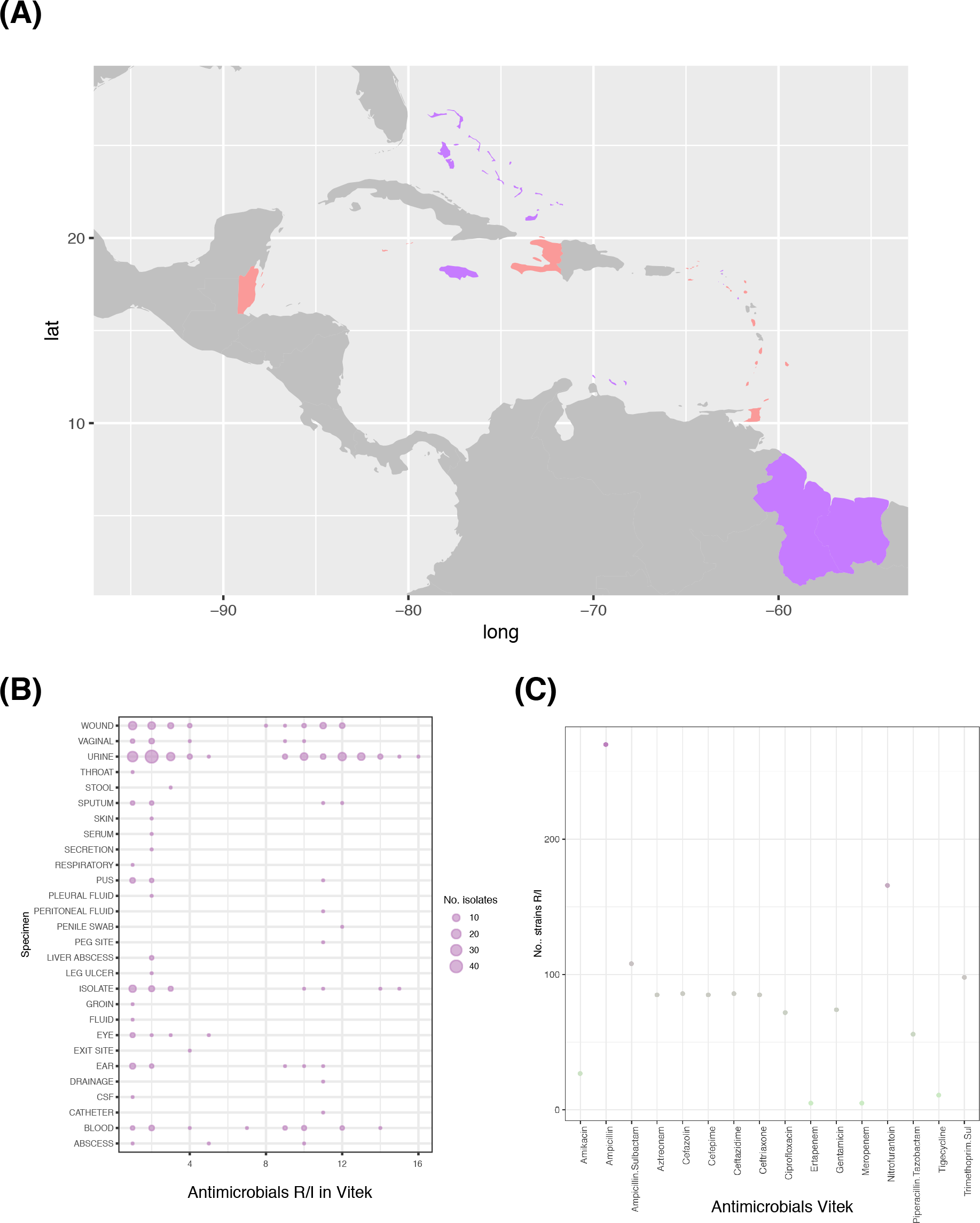
Characteristics of the collected isolates. **(A)** Map showing the CMS (violet), highlighting the CMS that contributed samples (red). **(B)** Vitek phenotypic resistance profiles of the analysed strains in the context of the diverse specimens the isolates derived from. **(C)** Number of strains R/I shown for each relevant antimicrobial measured.

Using whole-genome sequences, the Caribbean isolates were compared to a global collection designed to capture the *K. pneumoniae* species population diversity [5]. The *K. pneumoniae* population in the Caribbean show similar diversity when considering the O-antigen or capsule loci, as well as the range of molecular sequence types present in this region (Figure 3A), meaning that they are not comprised of only one or few widespread lineages in the Caribbean, but are representative of a genetically diverse established population. Our data shows a high diversity of isolates including all subspecies of the sequence complex (*K. pneumoniae*, *K. quasipneumoniae*, *K. variicola;* Figure 3B), and we note an expansion in the *K. quasipneumoniae* subsp. *similipneumoniae*, which has only recently been recognised as an important contributor to hospital isolates [15], as well as a relative enrichment in *K. quasipneumoniae* subsp. *quasipneumoniae*, whereas *K. variicola* seems underrepresented. Phylogenetic analysis clearly shows that the Caribbean isolates represent the broad *K. pneumoniae* diversity, as established in Holt et al 2015 (Figure 3C) [5].

**Figure 3:**
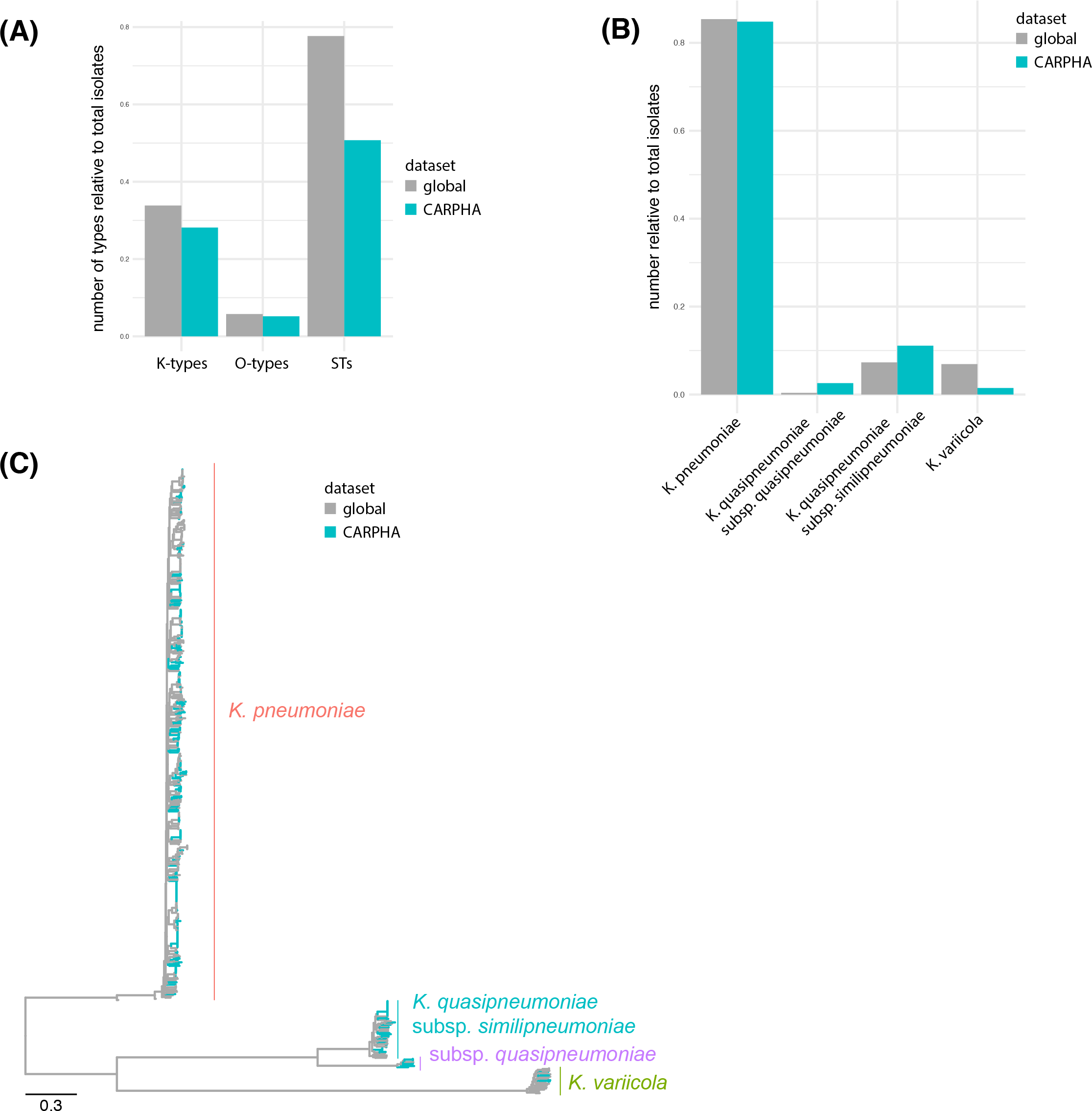
The Caribbean data in a global context. **(A)** Comparison of the diversity based on capsule (K), O-antigen (O) and sequence types (STs) compared with the global population study of Holt et al 2015. **(B)** Comparison of the three different species between the global collection and this study. **(C)** Phylogenetic analysis of the retrieved isolates demonstrates that they represent the global *K. pneumoniae* population as established by Holt et al. 2015[5].

There is little recent information available about how widespread AMR is in the region apart from single country reports[16][17][18], and no CARPHA member state had enrolled in the Global Antimicrobial Resistance Surveillance System (GLASS). However, the Caribbean is located between two hotspots of carbapenem-resistant *K. pneumoniae*, which are a recognised high risk in the U.S. (https://www.cdc.gov/hai/organisms/cre/index.html), and levels in South America are equally very high in several countries, with ESBL-resistance over 80% and over 25% carbapenem-resistant isolates (http://www.paho.org/data/index.php/en/mnu-topics/antimicrobial-resistance/320-klebsiella-spp.html). Our analysis of the genomic data shows a high number of acquired drug resistance genes present in the genomes of a considerable number of isolates and sequence types. The genotypic predictions of resistance largely match with their phenotypic resistance profiles (Figure 4). The majority of observed resistant isolates are clustered in several sequence types (STs); including ST11, ST15, ST29, ST392, and ST405 [19][20]. Also present at low numbers even in our limited number of samples, are high-risk clones such as ST258 [21]; although this isolate did not carry a carbapenemase gene. However, one isolate belonging to ST11, was found to carry carbapenemase KPC-1 and the AmpC cephalosporinase DHA-1 (Figure 4).

**Figure 4:**
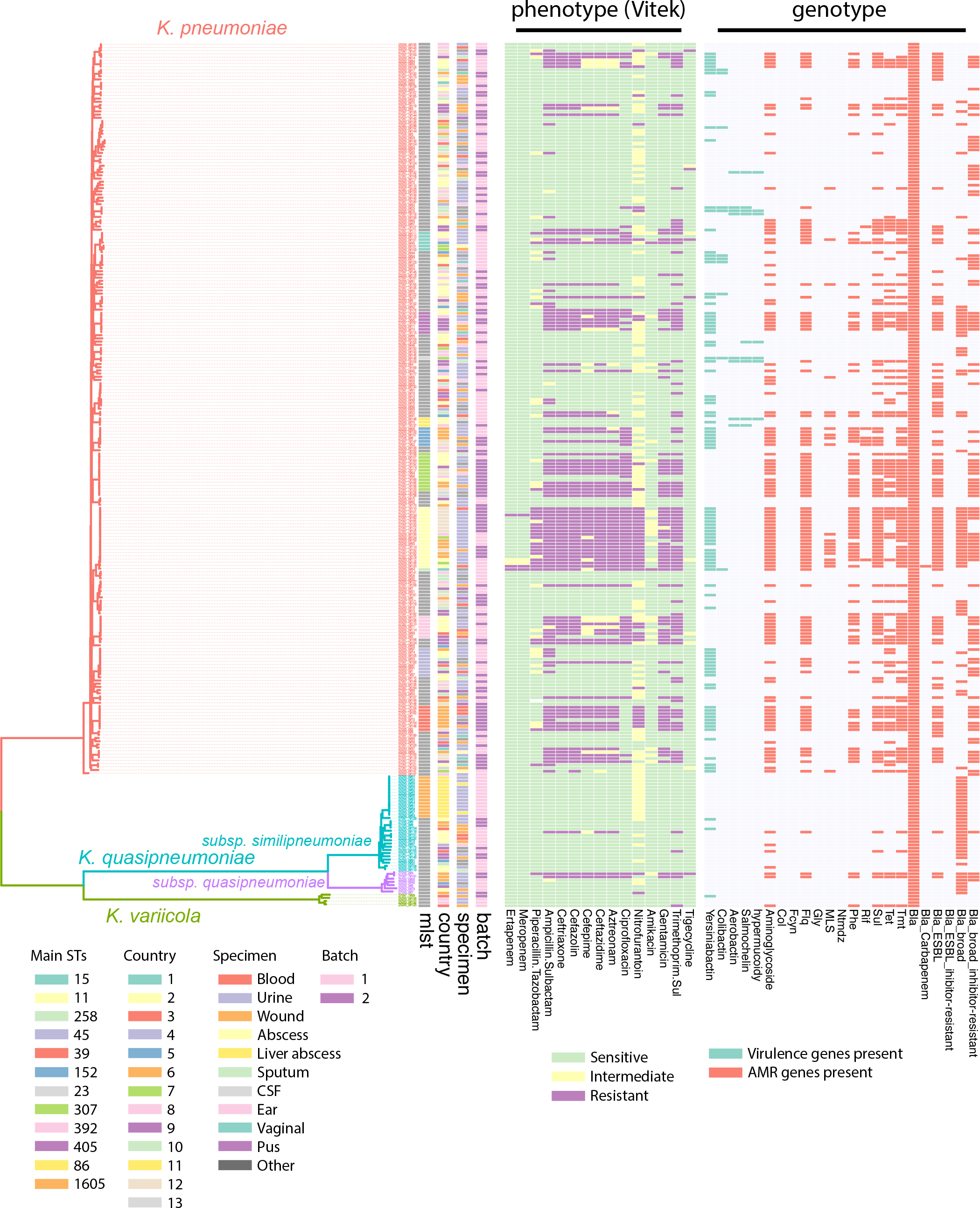
Whole-genome sequencing analysis reveals several high-risk clones with high antimicrobial resistance and hypervirulent lineages. The guidance tree is based on the core gene alignment as obtained by roary, and the colour strips represent, from left, the major sequence types as determined by MLST, the country-code of isolation and the specimen from which the isolate was obtained, and whether these were delivered in the first or second batch (early/late 2017). The heat maps represent the measured resistance phenotype as determined by the Vitek (green, sensitive; yellow, intermediate; violet, resistant), and the predicted resistance genes (red) and main virulence operons (green).

In addition to the main high-risk clones, with respect to antimicrobial resistances, we also noticed isolates belonging to hypervirulent lineages: an ST23 isolate, two isolates belonging to ST65 (AMR0288 and AMR0296), and ST86 [22][23][24][25][26]. Whilst the ST23 isolate was submitted as urine isolate (AMR0157), one of the ST86 isolate (17-02612) encoding a high number of virulence factors is derived from a fatal meningitis case, and the lineage is closely related to the previously reported strain from a case in Guadeloupe [26], whilst a second ST86 isolate (AMR0062) seems to miss/have lost the virulence plasmid (Figure 4).

Comparing the number of resistance and virulence determinants shows the typical split distribution with highly virulent and highly resistant strains, but the convergence between virulence and resistance cannot be observed in our limited sampling data, although all ingredients are present in the local gene pool (Figure 5A). Whilst the resistant isolates are strongly represented in blood and urine isolates (Figure 5B), the origins of highly virulent strains are more diverse and includes sterile sites such as CSF as expected (Figure 5B).

**Figure 5:**
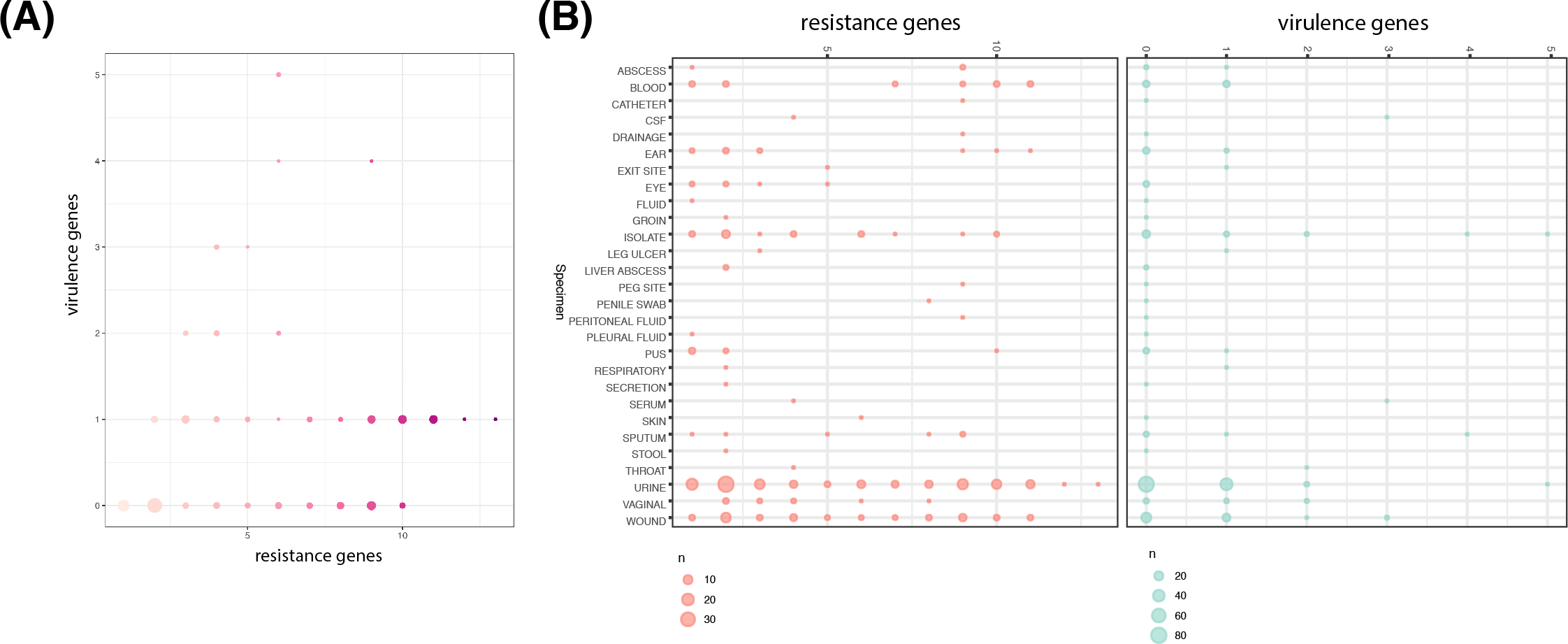
Distinct populations multidrug resistant and hypervirulent. **(A)** Comparison of the number of virulence and resistance determinants per strains shows bimodal distribution. **(B)** Comparison of the sample specimen, resistance (left panel) and virulence determinants (right panel).

## DISCUSSION

This study aimed to establish a genomic surveillance network across the Caribbean. Although this study was prematurely terminated it has provided important data. We show that several high-risk multidrug-resistant bacterial clones are present in clinical samples collected across the Caribbean. For *K. pneumoniae* these include sequence types ST258, ST11, ST15 and ST405. The diversity of the high-risk clones highlights that the risks from AMR are not limited to or described by the spread of a single high-risk lineage across different states or islands. Neither do we see a single plasmid or genetic island moving between relevant pathogens causing disease, but a large pool of diverse *K. pneumoniae* lineages and resistance elements distributed across region.

Given the limitations and the lack of a structured surveillance framework, we cannot conclude whether our data accurately reflects the true prevalence or the full extent of spread of these bacteria across this region. However, even given our limited sampling, there is a significant risk of rapid spread or ongoing, unnoticed epidemics of some of these high-risk clones, the presence and circulation of which will be hidden from view without using highly accurate approaches such as WGS.

Descriptions of infections caused by hypervirulent *K. pneumoniae* strains in the Caribbean are so far rare [26][27], but given the lack of surveillance, these observations might only represent the tip of the ice berg. The presence of high-risk multidrug resistant and hypervirulent strains, most of which are carried on mobile elements, also bears the further threat of the convergence to a multidrug resistant hypervirulent strain, as reported for e.g. ESBL-positive ST29 [28], KPC-positive ST11 acquiring hypervirulence features [24][29], or hypervirulent ST23 acquiring AmpC DAC-1 and ESBL enzymes [30]; all these components are part of the *K. pneumoniae* pool circulating in the Caribbean.

We argue that it is of crucial importance to continue the building of a systematic surveillance framework in the Caribbean, to fully assess the situation, provide informed guidelines for antimicrobial use and update these, as well as monitor high-risk clones and prevent outbreaks at their start.

## Acknowledgements

We thank the CARPHA member states public and hospital laboratories for their participation in this study. We thank the Wellcome Trust Sanger Institute Pathogen Informatics team for expert informatics support.

## Ethical statement

No ethics permission was required for the antibiotic point prevalence surveys as these were service reviews. All data was anonymised by the submitting hospitals.

## Author contributions

EH, RB and NRT conceptualised and designed the study. AMM sourced the isolates. AMM and KP identified and performed susceptibility testing, and EH analysed the data. EH, RB and NRT interpreted the data and wrote the manuscript.

## ABBREVIATIONS

CARPHA: Caribbean Public Health Agency
KPC: *K. pneumoniae* carbapenemase
ESBL: extended-spectrum beta-lactamase
SRA: sequence read archive
CMS: CARPHA member states
CSF: cerebrospinal fluid

